# The anti-apoptosis ubiquitin E3 ligase XIAP promotes autophagosome-lysosome fusion during autophagy

**DOI:** 10.1101/291294

**Authors:** Isabella Poetsch, Petra Ebner, Luiza Deszcz, Iris Bachtrog, Madlen Stephani, Carlos Gómez Díaz, Yasin Dagdas, Fumiyo Ikeda

## Abstract

The Inhibitor of Apoptosis Protein (IAP) family members are well-known endogenous regulators of apoptosis. Whether these proteins regulate other degradation pathways is unclear. Here, we discovered that the IAP member X-linked IAP (XIAP) is crucial for macroautophagy. Loss of XIAP in mouse and human cells inhibited starvation-induced degradation of LC3 proteins and an autophagy substrate p62. It also led to the accumulation of mature autophagosomes, suggesting that XIAP controls autophagic flux by mediating autolysosome formation. *Xiap*^*∆RING/∆RING*^ cells phenocopy the autophagy defects of *Xiap*^−/−^ cells, suggesting that the ubiquitinating activity mediated by the catalytic RING domain is critical for autophagic flux. We found that XIAP physically interacts with Syntaxin 17, a regulator of autophagosome-lysosome fusion. Syntaxin 17-positive mature autophagosomes positive accumulate in the cytoplasm of starved *Xiap*^−/−^ cells, suggesting that XIAP might regulate its dissociation from autophagosomes after fusion. XIAP selectively interacts with GABARAP among LC3 family members via the LIR-Docking Site (LDS). Together, our data suggest that XIAP-mediated ubiquitination regulates key autophagy regulators to promote autophagosome-lysosome fusion.

## Introduction

Autophagy is a critical cellular process that maintains cellular and tissue homeostasis by clearing autophagy cargos such as damaged organelles, protein aggregates and bacteria via the autolysosome (Feng, He et al., 2014, Jiang & Mizushima, 2014, Mizushima & Komatsu, 2011, Mizushima, Levine et al., 2008). Non-selective macroautophagy is initiated by starvation, which triggers the formation of double membrane phagophores around cargo. Phagophores become LC3-positive autophagosomes, which develop into Syntaxin 17-positive mature autophagosomes that fuse with lysosomes to form autolysosomes (Itakura, Kishi-Itakura et al., 2012, Mizushima, Yoshimori et al., 2011). Cargos are then hydrolyzed in autolysosomes by lysosomal enzymes (Shen & Mizushima, 2014). Previous studies have identified many autophagy regulators, including Autophagy-related (ATG) proteins, Unc-51 Like autophagy activating Kinase 1 (ULK1) and Beclin 1, most of which are critical for the initiation of autophagosome formation (Mizushima et al., 2011, Nakatogawa, Suzuki et al., 2009). More recently, multiple proteins important for autophagosome-lysosome fusion have been identified, including the Homotypic fusion and vacuole Protein Sorting (HOPS) tethering complex components, Pleckstrin Homology domain-containing family M member 1 (PLEKHM1), Rab7, EPG5, ATG14, Vesicle Associated Membrane Protein 8 (VAMP8), Soluble NSF Attachment Protein 29 (SNAP29) and the SNAP Receptor (SNARE) protein, Syntaxin 17 (Diao, Liu et al., 2015, Hyttinen, Niittykoski et al., 2013, Itakura et al., 2012, Itakura, Kishi et al., 2008, Jiang, Nishimura et al., 2014, Kummel & Ungermann, 2014, McEwan, Popovic et al., 2015, Wang, Miao et al., 2016).

A type of autophagy called selective autophagy targets ubiquitinated cargos, such as damaged mitochondria and aggregates as well as ubiquitin-coated bacteria (Khaminets, Behl et al., 2016, Shaid, Brandts et al., 2013), indicating that ubiquitination can play an important role in autophagy-dependent degradation. Some of the regulators of selective autophagy have been identified, especially those targeting damaged mitochondria (mitophagy), such as the ubiquitin E3 ligase Parkin, and PTEN-induced putative kinase 1 (PINK1) (Durcan & Fon, 2015, Harper, Ordureau et al., 2018, Nguyen, Padman et al., 2016, Pickrell & Youle, 2015, Yamano, Matsuda et al., 2016). However, only a handful of ubiquitin enzymes are known to regulate selective autophagy. Further, the roles for ubiquitin enzymes in macroautophagy are poorly understood.

In a recent study, we identified a ubiquitin conjugating E2 enzyme called BRUCE/Apollon as a new mammalian macroautophagy regulator, controlling autophagosome-lysosome fusion (Ebner, Poetsch et al., 2018). BRUCE is a member of the IAP family, and inhibits apoptosis by targeting apoptosis regulators Caspase 9 and SMAD for ubiquitin-mediated proteasome-dependent degradation (Hao, Sekine et al., 2004, Lotz, Pyrowolakis et al., 2004). The IAP family consists of 8 members, all of which contain a single or multiple Baculovirus IAP Repeat (BIR) domains, which bind to substrates as Caspase 9 and SMAD (Silke & Vucic, 2014). Among them, XIAP, cIAP1, cIAP2, ML-IAP and ILP-2 are RING-type ubiquitin E3 ligases (Middleton, Budhidarmo et al., 2014, Silke & Vucic, 2014). It is currently not known whether any of these IAP family members regulate macroautophagy.

XIAP is frequently overexpressed in cancer and contributes to tumor survival (de Almagro & Vucic, 2012, Silke & Vucic, 2014). XIAP has been also associated with autoimmune diseases, including Crohn’s disease (Schwerd, Pandey et al., 2017). Patients with Crohn’s disease often contain mutations in *Xiap* that localize to the C-terminal half of the coding region and would delete the Ubiquitin-Associated (UBA) and the catalytic RING domains (Schwerd, Pandey et al., 2016). These mutations are associated with defects in antibacterial autophagy in intestine (Schwerd et al., 2017). In cancer cells, XIAP is mainly considered as an inhibitor of autophagy, based on the observation of increased lipidated-LC3s by XIAP deficiency (Lin, Ghislat et al., 2015). It has also been shown that XIAP plays a role in mitophagy (Hamacher-Brady, Choe et al., 2014, Xie, Lin et al., 2017). Here, we demonstrate that XIAP, via its E3 ligase activity, positively regulates macroautophagy (hereafter referred to as autophagy) by promoting autophagosome-lysosome fusion.

## Results

### Loss of the inhibitor of apoptosis protein XIAP disrupts autophagic flux

We previously identified the inhibitor of apoptosis (IAP) protein BRUCE as a crucial positive regulator of autophagic flux (Ebner et al., 2018). Based on this finding and the reported crosstalk between apoptosis and autophagy (Galluzzi Vitale et al., 2018), we investigated whether other potent IAP family members, namely XIAP, cIAP1 and cIAP2, regulate autophagy. We induced autophagy via amino acid starvation and monitored autophagic flux by detecting the levels of the autophagy receptor p62 and of the autophagosome markers LC3B and GABA Type A Receptor-Associated Protein (GABARAP). We found that autophagic flux was unaffected in MEFs with a single knockout of cIAP1 or cIAP2, whereas double deficiency of cIAP1 and cIAP2 led to elevated levels of p62, LC3B and GABARAP in basal and starved conditions, suggesting that they have redundant roles in autophagy (Supplementary Fig. 1a, lane 1-3 vs. lane 10-12).

In contrast, depletion or loss of XIAP alone impaired autophagic flux under starved condition in MEFs, HCT116 cells, and human haploid cells (HAP1) (Fig. 1a, Supplementary Fig. 1b-g). Relative to controls, XIAP-knockout and knockdown-MEFs, XIAP-knockout HCT116 cells, and XIAP-knockout HAP1 cells displayed elevated levels of lipidated and cytosolic (non-lipidated) endogenous LC3B in basal as well as starved conditions, and increased levels of p62 (Fig. 1a lane 1-4 vs. lane 5-8, Supplementary Fig. 1c lane 1-4 vs. lane 5-8, Supplementary Fig. 1g lane 1-4 vs. lane 5-8 or 9-12). Unlike p62 and LC3B, XIAP protein levels were not altered by inducing autophagy via starvation, or by abrogating autophagy via loss of ATG5 (Fig. 1a and b), suggesting that XIAP is not an autophagy substrate.

**Figure 1.**
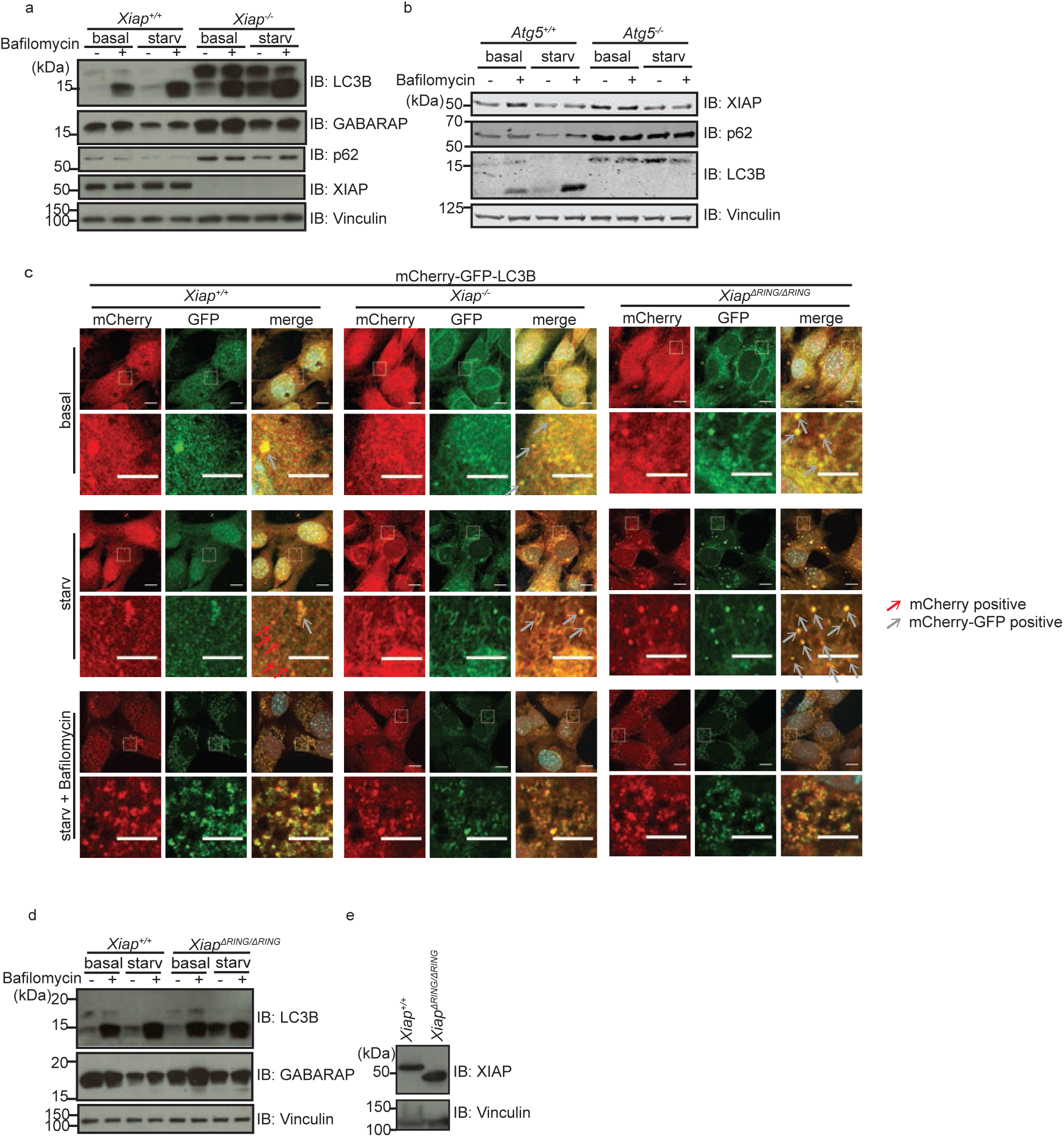
Degradation of LC3B, GABARAP and p62 is suppressed in *Xiap^−/−^* and *Xiap*^ΔRING/ΔRING^ MEFs by starvation-induced autophagy. **(a)** Levels of autophagy-related proteins in total cell extracts of XIAP wild type (*Xiap*^*+/+*^) and XIAP knockout (*Xiap*^−/−^) MEFs in different conditions were analyzed by immunoblotting. MEFs in basal or starved conditions with or without Bafilomycin A1 treatment (100nM, 2 hrs). Protein levels of LC3B, GABARAP, p62, XIAP are examined by using antibodies as indicated. Vinculin was monitored as loading control. **(b)** XIAP and p62 protein levels in *Atg5*^*+/+*^ and *Atg5*^−/−^ MEFs in basal and starved conditions with or without Bafilomycin A1 were examined by immunoblotting. Autophagy deficiency in *Atg5*^−/−^ MEFs was confirmed by levels of p62 and lipidation of LC3B. Vinculin was monitored as loading control. **(c)** Cellular localization of LC3B in *Xiap*^*+/+*^, *Xiap*^−/−^ and *XiapΔRING/ΔRINGΔ* MEFs stably expressing mCherry-EGFP-LC3B was analyzed by confocal microscopy. mCherry and GFP signals in MEFs in basal, or starved conditions with or without Bafilomycin A1 for 6 hours were examined. Scale bars, overview: 10µm, magnified view: 5µm. Gray and red arrows point to mCherry-GFP double positive and GFP-positive vesicles, respectively. **(d)** Autophagy-related proteins (LC3B and GABARAP) in total cell extracts of *Xiap*^*+/+*^ and XIAP ΔRING (*XiapΔRING/ΔRINGΔ*) MEFs were examined by immunoblotting. **(e)** Expression of WT XIAP and XIAP ΔRING proteins in *Xiap*^*+/+*^ and *XiapΔRING/ΔRINGΔ* MEFs was confirmed by immunoblotting using an anti-XIAP antibody. Vinculin was monitored as loading control.

To confirm the defect in autophagy flux, we monitored a stably-expressed tandem fluorescent LC3B fusion protein (mCherry-GFP-LC3B). Autophagosomes containing this pH-sensitive reporter protein stably maintain mCherry signal but lose the GFP signal upon fusion with lysosomes (Fig. 1c). Upon starvation, WT MEFs showed mCherry-positive-GFP-negative vesicles, whereas XIAP-knockout MEFs showed mCherry-GFP double positive vesicles, suggesting a defect in autophagic flux compared to WT cells (Fig. 1c). *Xiap*^−/−^ MEFs in basal and starved conditions displayed unaltered levels of proteins that are critical to initiate autophagosome formation, including ULK1, Beclin 1 and ATG4B, and of mRNAs encoding known autophagy regulators (Supplementary Fig. 1h-i), suggesting that defective autophagic flux in XIAP-deficient cells is not due to suppressed induction of known autophagy genes.

### Deletion of the XIAP RING domain impairs autophagic flux

XIAP is a well-described RING-type E3 ubiquitin ligase. To determine whether its ubiquitinating activity is required to promote autophagic flux, we evaluated a deletion mutant of XIAP that lacks the catalytic RING domain (XIAP-ΔRING). MEFs derived from XIAP-ΔRING mice (*Xiap*^*∆RING/∆RING*^ MEFs) phenocopied the autophagy defect observed in XIAP-deficient cells (Fig. 1c-e, Supplementary Fig. 1h and j). Relative to WT MEFs, *Xiap*^*∆RING/∆RING*^ MEFs displayed defective autophagy-dependent degradation of endogenous LC3B and GABARAP proteins (Fig. 1d lane 3 vs. lane 7 and Fig.1 e), in line with increased mCherry-GFP double positive LC3B vesicles upon starvation (Fig. 1c). Similar to XIAP knockout MEFs, *Xiap*^*∆RING/∆RING*^ MEFs showed no observable differences in proteins levels or mRNA levels of various known autophagy-related genes (Supplementary Fig. 1h and j).

### XIAP regulates autolysosome formation

Since autophagy flux was inhibited in XIAP-deficient cells and *Xiap*^*∆RING/∆RING*^ MEFs, we next examined whether this phenotype reflects disrupted autolysosome formation. To this end, we performed confocal microscopy to analyze autolysosomes via co-immunostaining for the endogenous lysosomal protein LAMP2 and the autophagosome membrane-conjugated protein LC3B in MEFs, in basal and starved conditions. We blocked autolysome turnover by treating cells with Bafilomycin A1, and found that starved *Xiap*^−/−^ and *Xiap*^*∆RING/∆RING*^ MEFs display a significant reduction in LAMP2-positive structures containing LC3B-positive autophagosomes (Fig. 2a and b). These data suggest that autolysosome formation is impaired upon loss of XIAP ubiquitinating activity, which would explain the defect in autophagic flux and the reduced degradation of endogenous autophagy substrate p62 and LC3 proteins (Fig. 1a, Supplementary Fig. 1c and g). In addition, we found that the average diameter of LAMP2-positive vesicles was significantly smaller in *Xiap*^−/−^ and *Xiap*^*∆RING/∆RING*^ MEFs (Fig. 2c).

**Figure 2.**
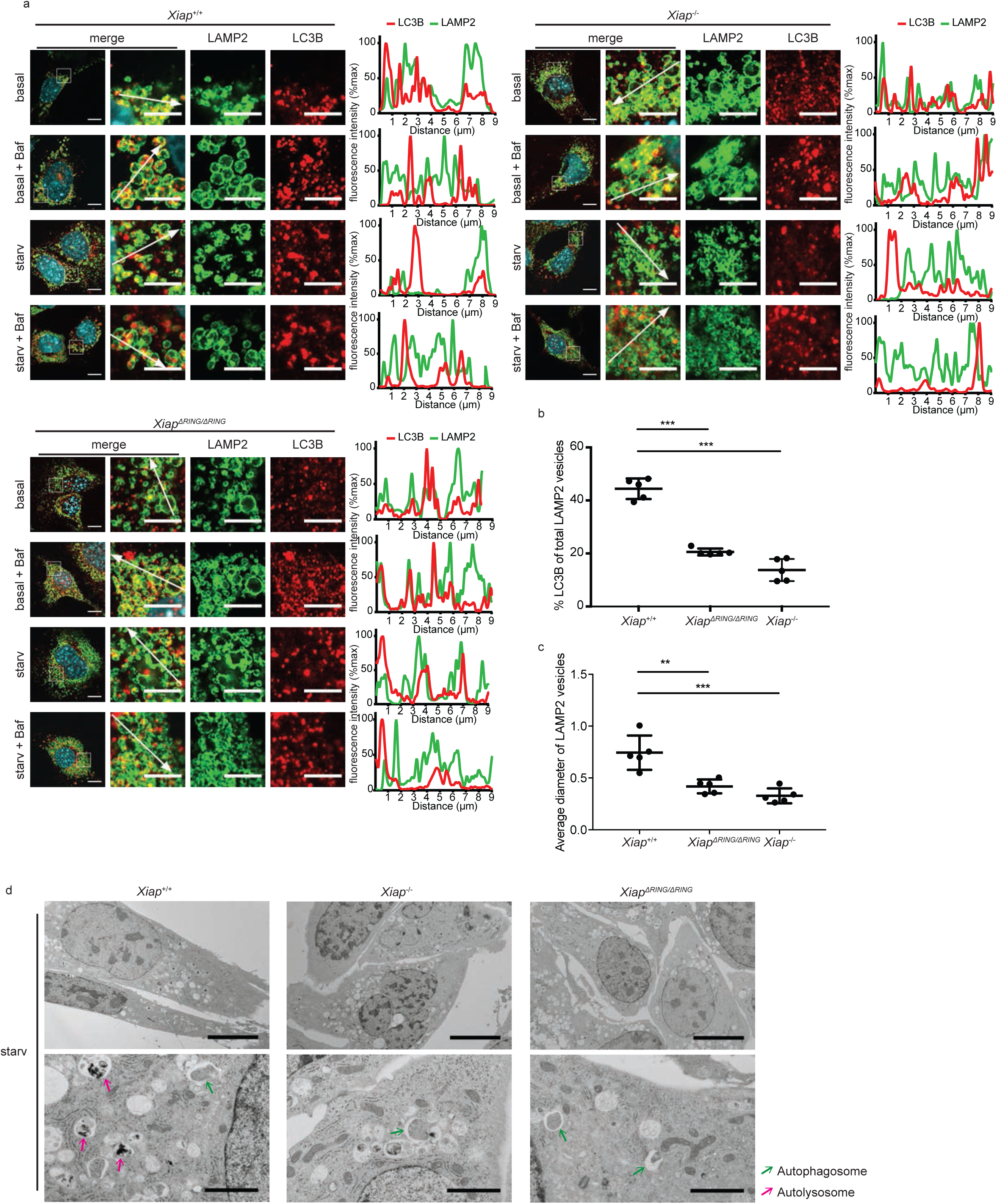
Autolysosome formation is defective in *Xiap*^−/−^ and *Xiap*^*∆RING/∆RING*^ MEFs. **(a)** Immunofluorescence staining of endogenous LC3B and LAMP2 in *Xiap*^*+/+*^, *Xiap*^−/−^ and *Xiap*^*ΔRING/ΔRING*^ MEFs. MEFs were starved for 2 hours and treated with Bafilomycin A1 (100nM) if indicated, and co-stained with anti-LC3B and anti-LAMP2 antibodies. Fluorescence intensity of each channel was blotted along the white arrow. Scale bars, overview: 10µm, magnified image: 5µm. Correlating changes in fluorescence intensity along regions marked with arrows are shown in line plots. **(b and c)** Quantification of (a) showing percentage of LAMP2-positive structures containing LC3B-positive aggregates (b) and average diameter of LAMP2 positive vesicles (c) in MEFs under starvation condition with Bafilomycin A1 treatment. Data are presented as dot plots with mean±SD (n=5; *p<0.05, **p<0.01, ***p<0.001). **(d)** Electron microscopic images of *Xiap*^*+/+*^, *Xiap*^*∆RING/∆RING*^ and *Xiap*^−/−^ MEFs starved for 2 h. Overview (top panels), scale bars, 10 µm; magnified view (bottom panels), scale bars are 2 µm. Green and pink arrows point to autophagosomes and autolysosomes, respectively.

We confirmed these observations by electron microscopy (Figure 2d, Supplementary Fig. 2a). Starvation induced the formation of autophagosomes in WT, *Xiap*^−/−^ and *Xiap*^*∆RING/∆RING*^ MEFs (Supplementary Fig. 2a, Fig. 2d). Although a substantial number of autolysosomes were visible in WT MEFs, only few were observed in *Xiap*^*-/-*^ and *Xiap*^*∆RING/∆RING*^ MEFs (Fig. 2d), pointing to a defect in the late steps of the autophagy pathway. Upon starvation and treatment with Bafilomycin A1, *Xiap*^−/−^ and *Xiap*^*∆RING/∆RING*^ MEFs displayed an increased number of LAMP2-positive structures relative to WT MEFs while a diameter of these vesicles was decreased (Supplementary Fig. 2b, Fig. 2c). We confirmed that *Xiap*^−/−^ and *Xiap*^*∆RING/∆RING*^ MEFs displayed unaltered levels of the major lysosomal proteins LAMP1 and LAMP2, and of the mRNAs encoding lysosomal biogenesis genes (Supplementary Fig. 2c-e). Together, these data suggest that autolysosome formation is defective in both *Xiap*^−/−^ and *Xiap*^*∆RING/∆RING*^ MEFs, and that XIAP promotes autolysosme formation through its catalytic RING domain.

### >Syntaxin 17-positive mature autophagosomes accumulate upon loss of XIAP ubiquitinating activity

Although *Xiap*^−/−^ and *Xiap*^*∆RING/∆RING*^ MEFs display a reduced number of autolysosomes, autophagosome formation appears to be intact. To clarify whether *Xiap*^−/−^ and *Xiap*^*∆RING/∆RING*^ MEFs accumulate mature autophagosomes, we monitored the subcellular distribution of Syntaxin 17 (STX17), which is recruited to mature autophagosomes and released upon autolysosome formation (Tsuboyama, Koyama-Honda et al., 2016). We generated WT, *Xiap*^−/−^ and *Xiap*^*∆RING/∆RING*^ MEF lines stably expressing GFP-STX17 and monitored its subcellular localization in different conditions. *Xiap*^−/−^ and *Xiap*^*∆RING/∆RING*^ MEFs displayed an increased number of mature GFP-STX17-positive autophagosomes compared to WT MEFs, under both basal and starved conditions (Fig. 3a), suggesting that autophagosomes mature but do not fuse with lysosomes in the absence of XIAP ubiquitinating activity. To further analyze the dissociation of STX17 from LC3B-positive autolysosomes, we performed immunostaining to detect endogenous LC3B in MEFs expressing GFP-STX17. Relative to WT MEFs, *Xiap*^−/−^ and *Xiap*^*∆RING/∆RING*^ MEFs showed increased co-localization of GFP-STX17 and LC3B in both basal and starved conditions, indicating that dissociation of GFP-STX17 was suppressed (Fig. 3b). Since the subcellular distribution of GFP-STX17 was affected in *Xiap*^−/−^ and *Xiap*^*∆RING/∆RING*^ MEFs, we next asked whether XIAP interacts with STX17. We found that STX17 interacts with GST-tagged XIAP in a pulldown assay (Figure 3c). Interestingly, other known autophagosome-lysosomal fusion proteins such as SNAP29, VPS16, ATG14, PLEKHM1, and the lysosomal protein LAMP1 did not interact with XIAP (Fig S3a-e).

**Figure 3.**
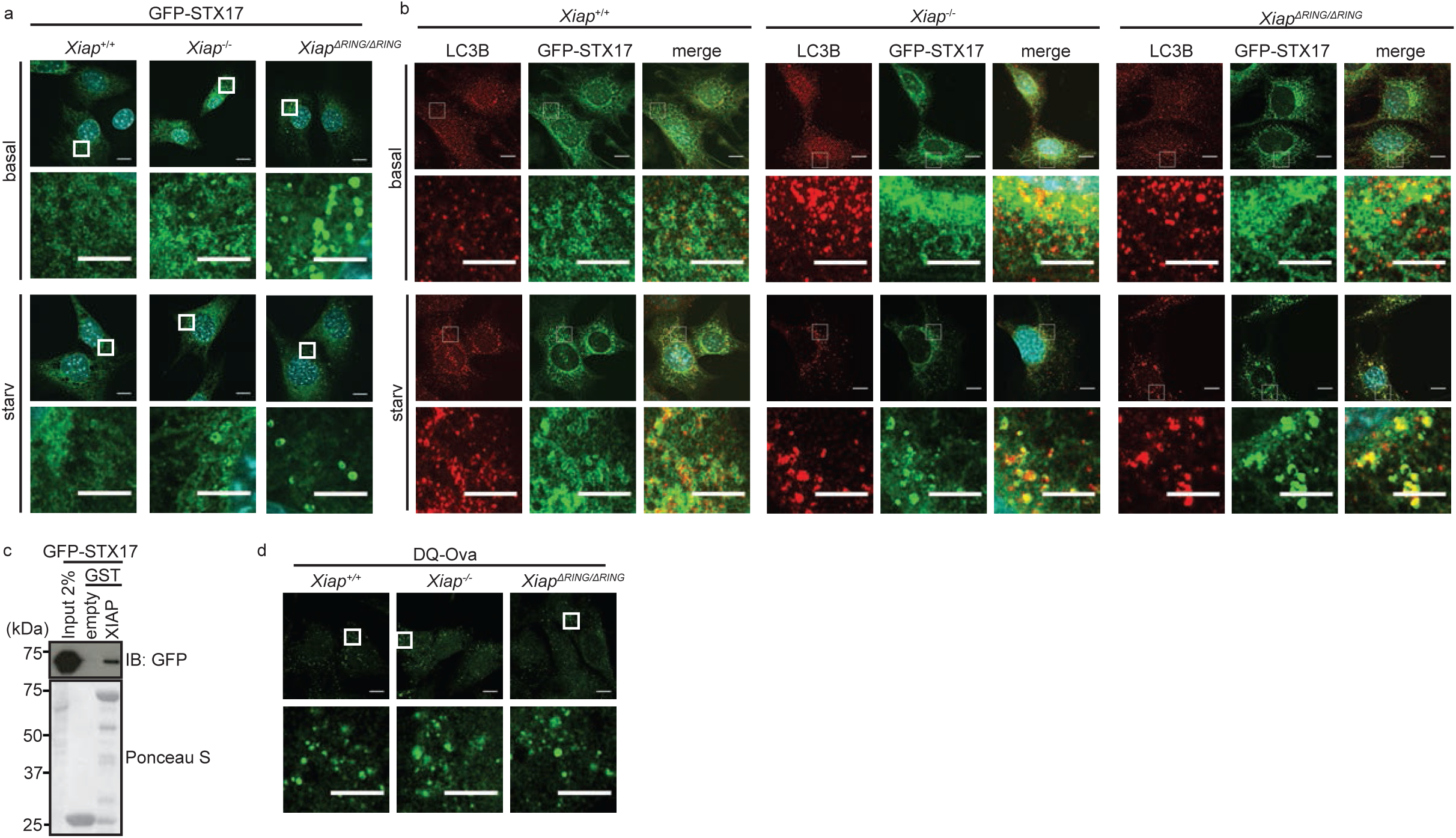
Syntaxin 17 (STX17)-positive mature autophagosome formation or the endocytosis pathway is not affected in *Xiap*^−/−^ and *Xiap*^*ΔRING/ΔRING*^ MEFs. **(a)** Confocal microscopic images of *Xiap*^*+/+*^, *Xiap*^−/−^ and *Xiap*^*ΔRING/ΔRING*^ MEFs stably expressing GFP-STX17 to visualize mature autophagosomes. MEFs in a basal condition or starved for 4h were fixed and stained with DAPI. Scale bars, overview: 10µm, magnified image: 5µm. **(b)** Confocal microscopy images of stably expressing GFP-STX17 and endogenous LC3B in *Xiap*^*+/+*^, *Xiap*^−/−^ and *Xiap*^*ΔRING/ΔRING*^ MEFs in basal or starved (2 hr) conditions. MEFs were fixed and stained with anti-LC3B antibody and DAPI. Scale bars, overview: 10µm, magnified image: 5µm. **(c)** Interaction between STX17 and GST-XIAP WT examined by GST-pulldown assays. Total cell extracts of HEK293T cells transiently expressing GFP-STX17 were incubated with agarose-immobilized GST-XIAP, pulled down, washed and examined by immunoblotting using an anti-GFP antibody. GST-empty was used as control and loading of GST-fusion proteins was examined by Ponceau S staining. **(d)** Functionality of endocytosis and lysosomal pathway was analyzed using DQ-Ovalbumin. Cells were grown in full medium or starved for 4.5 hr in DQ-Ovalbumin containing medium. Scale bars, overview: 10µm, magnified image: 5µm.

Some regulators of autophagosome-lysosome fusion also function in endocytosis (McEwan et al., 2014). To determine whether loss of XIAP causes defects in endocytosis or lysosomal activity, we evaluated fluorescence of the endocytic reporter DQ-Ovalbumin. DQ-ovalbumin is a self-quenched conjugate of ovalbumin that is endocytosed and exhibits fluorescence upon proteolytic cleavage in the lysosome (Lewis & Cobb, 2010). We observed similar levels of fluorescence in *Xiap*^*+/+*^, *Xiap*^−/−^ and *Xiap*^*∆RING/∆RING*^ MEFs treated with DQ-Ovalbumin, suggesting that lysosomal activity is intact in the absence of XIAP (Fig. 3d). Together, these data suggest that the RING domain of XIAP is required for proper autophagic flux, but not for receptor-mediated endocytosis.

### XIAP selectively interacts and colocalizes with the ATG8 family members GABARAP and GABARAPL1

To further understand how XIAP promotes autophagy flux, we examined potential physical interactions between XIAP and autophagosomal proteins, the LC3 family members. We performed GST-pulldown assays with a collection of XIAP mutants and mammalian LC3 proteins (Fig. 4a-c, Supplementary Fig. 4a-j). We found that, among six ATG8 family members, GST-GABARAP and GST-GABARAPL1 selectively interacted with HA-tagged WT XIAP from total cell extract of transiently transfected HEK293T cells (Fig. 4b). This interaction was independent of the XIAP RING domain (Fig. 4c). All of the XIAP mutants we examined, except HA-XIAP-BIR3, interacted selectively with GST-GABARAP or GST-GABARAPL1 (Fig. 4a, Supplementary Fig. 4a-i), suggesting that there are multiple interaction sites within XIAP. Recombinant WT XIAP protein interacted equally well with GABARAP and GABARAPL1, pointing to direct interactions (Supplementary Fig. 4j). We also detected weak interactions between recombinant XIAP and di-, tri- or tetra-ubiquitins (Supplementary Fig. 4j).

**Figure 4.**
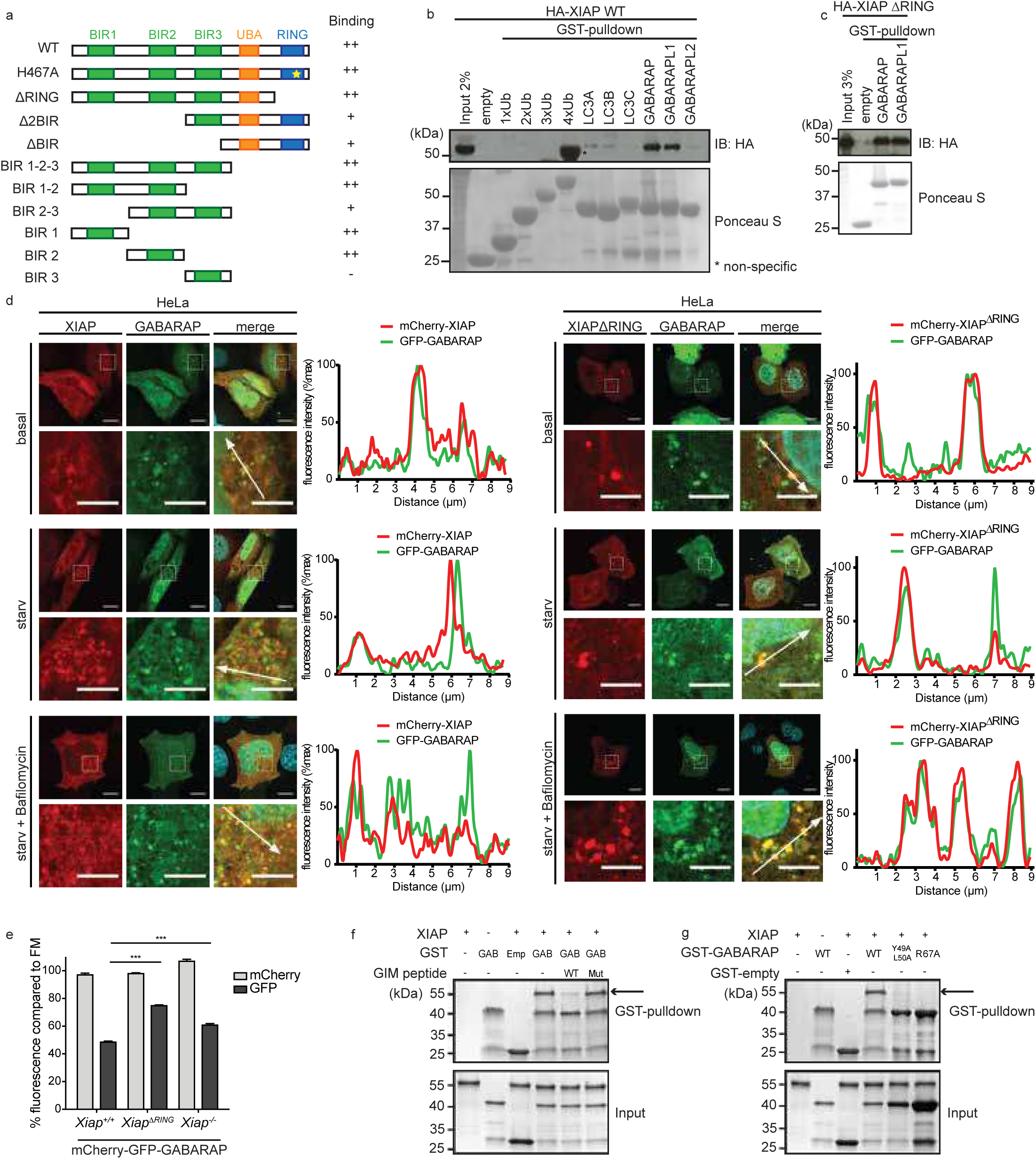
XIAP interacts and colocalize with GABARAP, and regulates GABARAP-positive autophagosome turnover. **(a)** A schematic of XIAP mutants used in the interaction assays with LC3 proteins. **(b and c)** Interaction of XIAP wild type (WT) or XIAP ΔRING and mammalian LC3 proteins and ubiquitins examined by GST-pulldown assays. Total cell extracts of HEK293T cells transiently expressing HA-tagged WT (b) or ΔRING (c) was incubated with GST-ubiquitin (monomer to tetramer) or GST-ATG8 proteins immobilized with glutathione sepharose beads. Interaction was detected by immunoblotting using anti-HA antibody and loading of GST proteins were examined by Ponceau S staining. **(d)** Confocal microscopy image of HeLa cells transiently expressing mCherry-XIAP and GFP-GABARAP or mCherry-XIAPΔRING and GFP-GABARAP. Cells were grown in basal conditions or starved for 2h with or without Bafilomycin A1 (100nM). Fluorescence intensity of each channel was blotted along the white arrow. Scale bars, 10µm and 5µm in magnified images. **(e)** Degradation of mCherry-EGFP-GABARAP upon starvation was analyzed in *Xiap*^+/+^, *Xiap*^−/−^ and *Xiap*^ΔRING/ΔRING^ MEFs by flow cytometry. mCherry-EGFP-GABARAP stably expressing MEFs were starved 6 hours and mean fluorescence signal was compared to basal conditions. Data are presented as mean±SD (n=3; ***p<0.001). Representative data are shown from three independent experiments. **(f)** Peptide competitions assays for a direct interaction between recombinant XIAP and GST-GABARAP (GAB) proteins. Interaction was determined by GST-pulldown assay with or without GABARAP-interaction motif (GIM)-peptides (WT or a mutant), followed by SDS-PAGE and Coomassie staining. GST-empty (Emp) was used as control. An arrow indicates bands corresponding to XIAP. **(g)** Direct interaction between XIAP and GABARAP (WT, Y49A/L50A or R67A) examined by GST-pulldown assay as in (f). An arrow indicates bands corresponding to XIAP.

Typically, LC3 interacting proteins contain a short “LC3 interacting region” (LIR) that is docked into a conserved surface on LC3 proteins, the so-called “LIR docking site” (LDS) (Kirkin, Lamark et al., 2009, Noda, Kumeta et al., 2008). To test whether the interaction between XIAP and GABARAP is based on binding between LIR-LDS motifs, we performed a peptide competition assay (Fig. 4f). We found that the interaction between recombinant XIAP and GABARAP was inhibited by the addition of WT LIR peptides, but not mutant LIR peptides, suggesting that XIAP contains a canonical LIR motif (Behrends, Sowa et al., 2010). Furthermore, mutations within the GABARAP-LDS (Y49A L50A, R67A) abrogate the interaction with XIAP (Fig. 4g), supporting that the XIAP-GABARAP interaction mode is mediated by LIR-LDS binding.

In line with the biochemical interaction detected between XIAP and GABARAP, we also observed co-localization of mCherry-XIAP and GFP-GABARAP in transiently transfected HeLa cells (Fig. 4d). Compared to XIAP-WT protein, XIAP-ΔRING proteins formed larger cytosolic aggregates that largely colocalized with GFP-tagged GABARAP (Fig. 4d). We found that starvation-induced quenching of the GFP signal of fluorescent mCherry-GFP-GABARAP is reduced in *Xiap*^*-/-*^ and *Xiap*^*∆RING/∆RING*^ MEFs relative to WT MEFs (Fig. 4e). Since WT XIAP and XIAP-∆RING interact equally well with GABARAP, these data suggest that the XIAP-GABARAP interaction alone is not sufficient for XIAP to promote autophagy, and that the XIAP RING domain is required.

## Discussion

Our work reveals that XIAP, specifically its RING domain, is required for a late step in the autophagy pathway but is dispensable for the formation of mature autophagosomes. Our data further suggest that efficient autophagosome-lysosome fusion is impaired upon loss of XIAP ubiquitinating activity, given that we observed accumulation of mature autophagosomes positive for LC3B or GABARAP, suppressed dissociation of STX17, and reduced formation of autolysosomes.

XIAP is a well-known cell death inhibitor. As a ubiquitin E3 ligase, XIAP ubiquitinates apoptosis regulators and targets them for proteasomal degradation (de Almagro & Vucic, 2012, Silke & Vucic, 2014). IAP genes are important for tumor survival and development; mutations, amplifications and chromosomal translocations of IAP genes are associated with various malignancies (Fulda & Vucic, 2012). Once cancer is established in humans, it is important to inhibit autophagy to suppress cancer (Jiang & Mizushima, 2014, White, Karp et al., 2010). Our data suggest that inhibiting XIAP would not only promote apoptosis but also suppress autophagy, both of which might be beneficial for cancer treatment. XIAP has been suggested to negatively regulate autophagy in cancer cells (Huang, Wu et al., 2013), whereas other studies suggest that XIAP is a positive regulator of autophagy (Huang, Wu et al., 2013, Lin et al., 2015). Moreover, a recent study has shown that Crohn’s disease patients acquire XIAP gene mutations (Schwerd et al., 2016). The autophagy gene *ATG16L1* is also mutated in some Crohn’s disease patients (Bhattacharya & Eissa, 2013, Fujita, Saitoh et al., 2009). These data suggest that inhibition of autophagy may contribute to Crohn’s disease.

Interestingly, unlike XIAP, single knockout of cIAP1 or cIAP2 was not sufficient to disrupt autophagy. However, MEFs with a double knockout of cIAP1 and cIAP2 showed a clear accumulation of LC3B, GABARAP and p62 upon starvation, suggesting that cIAP1 and cIAP2 redundantly promote autophagy flux. We found that defects in autophagy seemed more severe in XIAP-ΔRING MEFs than in XIAP knockout MEFs, especially in degradation of GABARAP-positive autophagosomes and dissociation of Syntaxin 17 from LC3-positive autophagosomes in starved cells. Because the RING domain contains the active site for ubiquitin E3 ligase catalysis, we expect that XIAP regulates autophagy regulators by ubiquitination. However, because deletion of the RING domain may abrogate additional functions of XIAP beyond its ubiquitinating activity, it remains possible that other potential functions of XIAP are required for autophagosome-lysosome fusion. XIAP is known to ubiquitinate itself with Lys48-linked ubiquitin chains, triggering its own proteasomal degradation (Sekine, Takubo et al., 2008, Yang, Fang et al., 2000). We did not observe changes in XIAP protein levels upon starvation of WT mammalian cells, or in autophagy-deficient ATG5 knockout MEFs, suggesting that the XIAP catalytic activity may not be directly regulated by autophagy induction. However, as deletion of the RING domain is sufficient to trigger the autophagy defect in MEFs, it could be that XIAP ubiquitinates only specific autophagy regulators in this context, for example those at the autophagosomes.

Regulators of autophagosome-lysosome fusion, like PLEKHM1, often also play a role in the endocytosis pathway (McEwan et al., 2015). However, in the case of XIAP, the endocytosis pathway was intact determined by DQ-Ovalbumin internalization assays. DQ-Ovalbumin fluoresces only when it reaches lysosomes after endocytosis. We observed similar levels of fluorescence in *Xiap*^*+/+*^, *Xiap*^−/−^ and *Xiap*^*∆RING/∆RING*^ MEFs treated with DQ-Ovalbumin. XIAP interacts selectively with Syntaxin 17 among the important proteins for autophagosome-lysosome fusion that we tested (SNAP29, ATG14, VPS16, and PLEKHM1). Syntaxin 17 is specifically localized at mature autophagosomes, and may underlie the specificity of XIAP for autophagosome-lysosome fusion.

Compared to the steps that initiate autophagosome formation, the mechanistic understanding of later steps in autophagy, particularly autophagosome-lysosome fusion, remains unresolved. The discovery that autophagosome-lysosome fusion is regulated by the autophagosome-specific SNARE protein Syntaxin 17 advanced the field substantially (Itakura et al., 2012). Syntaxin 17 dissociates from autophagosomes after fusion with the lysosome (Itakura et al., 2012, Jiang et al., 2014, Uematsu, Nishimura et al., 2017), but the mechanisms that regulate its dissociation are not known. We found that Syntaxin 17-LC3B double positive vesicles accumulate in starved MEFs upon loss of XIAP, suggesting that XIAP may regulate the removal of Syntaxin 17 from the mature autophagogomes. Alternatively, this phenotype might reflect the disrupted regulation of autophagosome-lysosome fusion upon loss of XIAP. Regardless, it is clear that XIAP promotes autophagosome-lysosome fusion and a further understanding of its function will deepen our understanding of the mechanisms that regulate this process.

## Materials and methods

### Plasmids

pEBB-HA-XIAP FL (aa1-497) and mutants (XIAP H467A, XIAPΔRING (aa 1-449), BIR1-2-3 (aa 1-350), ΔBIR (aa 331-497), Δ2BIR (aa262-497), BIR3 (aa 262-350), BIR2 (aa 124-261), BIR1 (aa 1-123), BIR2-3 (aa 124-350), BIR1-2 (aa 1-261)) were a gift from Colin Ducket (Addgene #25674, #25675, #25676, #25686, #25687, #25688, #25689, #25690, #25691, #25692, #25693) (Lewis, Burstein et al., 2004). GST-tagged human MAP1LC3A, human MAP1LC3B, human MAP1LC3C, human GABARAP, mouse GABARAPL1 and human GABARAPL2 in pETM30 vector were kindly provided by Felix Randow (von Muhlinen, Akutsu et al., 2012). pEGFPC1-GABARAP, pmCherryC1-GABARAP, pmCherryC1-GABARAPL1, pBabe-puro-mCherry-EGFP-GABARAP, pBABE-puro-mCherry-EGFP-GABARAPL1, pmCherryC1-XIAP, pmCherryC1-XIAPΔRING, and GST-tagged human XIAP were generated using a standard subcloning method. pCMV-Gag-Pol was used as a helper plasmid for retrovirus production, and shRNA against mouse XIAP (TTTTGTACTATATACATCTTAA) were cloned into pRSF91-SFFV-TagBFP-mirE-PGK-Neo-WPRE as previously described (Ebner et al., 2018, Fellmann, Hoffmann et al., 2013). pGex4T1-human ubiquitin 1x, 2x, 3x, and 4x were described elsewhere (Ikeda, Deribe et al., 2011). pBabe-puro mCherry-EGFP-LC3B was a gift from Jayanta Debnath (Addgene plasmid # 22418) (N’Diaye, Kajihara et al., 2009) and LAMP1-mGFP was a gift from Esteban Dell’Angelica (Addgene plasmid # 34831) (Falcon-Perez, Nazarian et al., 2005). pMRXIP GFP-STX17 was a gift from Noboru Mizushima (Addgene plasmid #45909) (Itakura et al., 2012).

### Antibodies

Anti-XIAP (BD, 610716), anti-Ubiquitin (P4D1, Santa Cruz, sc-8017), anti-HA.11 (Covance, MMS-101P), anti-Alpha-Tubulin (Abcam, ab15246), anti-Vinculin (Sigma-Aldrich, V9131), anti-LC3 (Nano Tools, 0260-100/LC3-2G6), anti-GABARAP (E1J4E, Cell Signalling, 13733), anti-GABARAPL1 (Abcam, ab86497), anti-P62/SQSTM1 (MBL, PM045), anti-ATG4B (Cell signaling #5299), anti-LAMP1 (Abcam, ab24170), anti-LAMP2 (Abcam, ab13524), anti-GST (Covance MMS-112P), Anti-ULK1 (Santa Cruz, sc-33182) and Anti-Beclin1 (Cell Signaling, 3738) antibodies were purchased and used according to the manufacturer’s recommendations.

### Cell lines

*Xiap*^*+/+*^, *Xiap*^−/−^, *Xiap*^*ΔRING/ΔRING*^ mouse embryonic fibroblasts (MEFs) were kind gifts from Philipp J. Jost (Technical University Munich, Germany) and Ulrike Resch (Med. Univ. Vienna, Austria) (Schile, Garcia-Fernandez et al., 2008, Yabal, Muller et al., 2014). cIAP1, cIAP2 knockout and double knockout MEFs were kindly provided by W. Wei-Lynn Wong (University of Zurich). MEFs, HeLa Kyoto cells (kindly provided by Daniel Gerlich, IMBA, Vienna, Austria), Platinum-E (Plat-E, Ecotropic) cells, Platinum-A (Plat-A, Amphotropic) cells and Human Embryonic Kidney (HEK) 293T cells (ATCC) were maintained at 37°C in 5% CO2, in Dulbecco’s Modified Eagle Medium High Glucose (Sigma) supplemented with 10% Fetal calf serum (Thermo Fisher Scientific), 100 U/ml penicillin-streptomycin (Sigma) and 2 mM L-glutamine (Sigma-Aldrich). HAP1 cells were purchased from Haplogen/Horizon Genomics and were maintained in Iscove’s Modified Dulbecco’s Medium (IMDM, Life technologies), supplemented with 10% fetal calf serum and 100 U/ml penicillin-streptomycin. *XIAP*^*y/+*^, *XIAP*^*y*/-^ HCT116 cells were kindly provided by Philipp J. Jost (Yabal et al., 2014) and maintained in McCoy’s 5A medium modified, with L-glutamine and sodium bicarbonate, supplemented with 10% fetal calf serum and 100U/ml penicillin-streptomycin. All the cell lines used in this study were tested for mycoplasma and confirmed to be negative.

### Retroviral infection

A method for retrovirus production is described elsewhere (Ikeda, Hecker et al., 2007). Briefly, retroviral plasmid and helper plasmid were transfected in packaging cell lines (Plat-E and Plat-A respectively), using a standard calcium-phosphate transfection protocol. 48 h post transfection, filtered condition media containing retroviral particles supplemented with 4 µg/ml polybrene (Sigma, H9268) were used to infect MEFs. To obtain stable cell lines, cells were selected using 1.5 mg/ml G418 (Gibco, 108321-42-2) or 4 µg/ml puromycin (Lactan GmbH, 240.3).

### Transfection

Cells were transfected by using GeneJuice or by calcium phosphate method. For GeneJuice, cells were seeded in 6-well-plates at 70% confluency. Per well, 100μl DMEM was mixed with 3μl GeneJuice and incubated 5 minutes at room temperature. DMEM/GeneJuice mixture was added on 1μg DNA, mixed and incubated for 15 minutes at room temperature. Subsequently, the mixture was added dropwise on the cells. For calcium phosphate transfection method, platinum-E or Platinum-A cells were seeded at 70% confluency. Solution A (5μg plasmid DNA, 2μg pCMV-Gag-Pol helper plasmid, 250μM CaCl_2_) and solution B (2x HBS [280mM NaCl, 50mM HEPES, 1.5mM Na_2_HPO_4_, 12mM Dextrose, 10mM KCl, pH 7.02]) were prepared. Solution B was added dropwise to solution A at a 1:1 ratio while blowing air through the solution to mediate immediate mixing. The mixture was incubated 15 minutes at room temperature and subsequently added dropwise on the cells. Cells were incubated for 48 hrs before harvesting.

### Generation of XIAP-deficient HAP1 by CRISPR-Cas9

Methods were described elsewhere (Ebner et al., 2018). In short, Design of small guide RNAs (sgRNA) was performed using the online CRISPR Design Tool from Zhang Lab (http://crispr.mit.edu/). sgRNAs (*gXIAP* #1 TTGGACCGAGCCGATCGCCG, *gXIAP* #2 ATGACAACTAAAGCACCGCA) were cloned into pSpCas9(BB)-2A-GFP(PX458) (a gift from Feng Zhang (Addgene plasmid # 48138). HAP1 cells were transfected using GeneJuice transfection reagent (VWR International) and FACS sorted for GFP-positive cells 48 h post transfection. A pooled cell clone was used for subsequent experiments and compared to a mock transfected (pSpCas9(BB)-2A-GFP empty vector) pool.

### Flow cytometry

MEFs stably expressing mCherry-EGFP-LC3B or mCherry-EFGP-GABARAP were seeded in triplicates and starved for 6h in EBSS (Invitrogen, #24010043) (N’Diaye et al., 2014). Subsequently, cells were trypsinized (Trypsin-EDTA (0.05%), phenol red Gibco, 25300054), centrifuged (1000rpm, 5min) and resuspended in 300μl PBS. Cells were analyzed using FACS LSR Fortessa. For the analysis of basal fluorescence levels, the mean of the relative fluorescence units was displayed. To display autophagy induction, the fluorescence level in full medium or DMSO control condition was set to 100% for every cell line and the remaining fluorescence signal after autophagy induction was calculated accordingly.

### Fluorescence microscopy

Cells were seeded on cover slips at low density. On the following day, cells were starved in EBSS, and if indicated treated with 100 nM Bafilomycin A1 (T Yoshimori, 1991) (Enzo Life Sciences, BML-CM110-0100) or equal amounts of DMSO. Cells expressing fluorescently tagged proteins were either fixed for 15 min at room temperature in 4% paraformaldehyde, or fixed in MeOH for 10min at −20°C, washed and subsequently transferred to mounting medium (VECTASHIELD, Szabo-Scandic, VECH-1000 (no DAPI) or VECH-1200 (containing DAPI)). For staining of endogenous proteins, cells were fixed in cold methanol for 10 min at −20°C, re-hydrated in cold PBS on ice, followed by 3 washes. Cover slips were blocked in 5% BSA in PBS for 1 h at room temperature. Primary antibody was diluted in blocking solution according to the manufacturer’s recommendations and incubated overnight on cover slips. Samples were washed in PBS and incubated in secondary antibody, Alexa Fluor 568 goat anti-mouse IgG (H+L) (Invitrogen, A11031), or Alexa Fluor 488 goat anti-rat IgG (H+L) (Invitrogen, A11006), diluted 1:1000 in blocking solution for 1 h at room temperature. Samples were washed, and cover slips were transferred to mounting medium. Samples were imaged by confocal microscopy or Airy scan on LSM780 or LSM880 Axio Observer (Zeiss). For the analysis of LAMP2 positive vesicles, the diameter, number and distance to the nucleus was manually quantified using Fiji software. For the quantification of LC3B containing LAMP2 vesicles, the number of strongly stained LC3B aggregates was counted manually using Fiji.

### Electron microscopic analysis

MEFs were grown on 12 mm Aclar plastic discs and fixed for 1 hour in 2.5% glutaraldehyde in 0.1 M sodium phosphate buffer, pH 7.4. Samples were then rinsed with the same buffer, subsequently fixed in 1% osmium tetroxide in ddH_2_O, dehydrated in a graded series of ethanol and embedded in Agar 100 resin. 70-nm sections were cut parallel to the substrate and post-stained with 2% uranyl acetate and Reynolds lead citrate. Sections were examined with an FEI Morgagni 268D (FEI, Eindhoven, The Netherlands) operated at 80 kV. Images were acquired using an 11-megapixel Morada CCD camera (Olympus-SIS).

### DQ-Ovalbumin endocytosis assay

To monitor changes in lysosomal enzymatic activity, cells were grown in full medium and starved for 4.5h with or without Bafilomycin A1 treatment (100nM). In the last 2h of starvation and treatment, medium was changed to 10µg/ml DQ-Ovalbumin (Thermo Fisher Scientific, D12053) containing medium. Cells on cover slips were washed, fixed for 15 min with 4% paraformaldehyde-PBS and transferred to mounting medium containing DAPI. Samples were analyzed using LSM880 Axio Observer (Zeiss).

### Immunoblotting

The method is described elsewhere (Ikeda et al., 2011). Briefly, cells were lysed with chilled lysis buffer (50mM HEPES, 150mM NaCl, 1mM EDTA, 1mM EGTA, 25mM NaF, 10 mM ZnCl_2_, 10% glycerol, 1% Triton X-100, 20 mM NEM and Complete protease inhibitors) and total cell lysates were separated by SDS-PAGE, and transferred to nitrocellulose (GE Healthcare, Little Chalfont, UK) or to PVDF membrane (Millipore, ISEQ00010). Membranes were blocked in 5 % BSA-TBS blocking solution (150mM NaCl, 50mM Tris, 0.1%Na-azide, 5% BSA, phenol red, pH7.5-7.7) and blotted with indicated antibodies in blocking solution at 4 °C overnight. Secondary antibodies were used according to the manufacturer’s recommendations: goat anti-mouse HRP (BioRad, 170-6516), goat anti-rabbit HRP (Dako, P0448) and goat anti-rat IgG HRP (Southern Biotech, 3050-05). Western Blotting Luminol Reagent (Santa Cruz) and High performance chemiluminescence films (GE Healthcare, Little Chalfont, UK) were used. Where appropriate, Ponceau S staining was used to visualize transferred proteins on the membranes.

### GST-fusion protein purification

A method for GST-protein purification and pulldown assay are described elsewhere (Ikeda et al., 2011). Briefly, transformed *Escherichia coli* BL21 (DE3) were grown overnight at 37°C. 5ml overnight culture was added to 400ml fresh LB medium until an OD_600_ of 0.35-0.6 was reached. Expression of GST-fusion proteins was induced with 0.1mM IPTG and grown overnight at 16°C. The culture was centrifuged (15 minutes, 6000g, 4°C, rotor JA-10) and the pellet was washed in PBS, resuspended in GST-Buffer (20mM Tris, pH8.0, 100mM NaCl, 1mM PMSF) and subsequently sonicated and lysed in 0.5% Triton X-100. Lysate was cleared by centrifugation (20 minutes, 10 000g, 4°C, rotor JA-30.50) and incubated with Glutathione Sepharose 4B agarose beads (GE Healthcare) 1 hour rotating at 4°C. Subsequently, beads were washed in GST-buffer 2 (0.1M Tris, pH7.8, 0.5M NaCl, 1mM PMSF, 0.1% β-mercaptoethanol) and resuspended in 800μl GST-buffer 3 (20mM Tris, 0.1% NaN_3_).

### XIAP purification

A method for protein purification is described elsewhere (Ikeda et al., 2011). Briefly, transformed *Escherichia coli* BL-21 strain (DE3) was grown at 37°C in 12l LB medium supplemented with 100μM ZnCl2 for ~2h until an OD_600_ of 0.4 was reached. The incubation temperature was reduced to 25°C and cells were grown to reach OD_600_ of 0.8. Expression of GST-XIAP was induced by addition of 100μM IPTG and cells were grown for 5h. Cells were centrifuged, resuspended in 100ml suspension buffer (100mM HEPES, 500mM NaCl, pH7.4, 3000U DNase I recombinant (Roche), 6 tablets cOmplete mini, EDTA-free proteinase inhibitor cocktail), 0.5ml PMSF (100mM in EtOH, Sigma Aldrich) was added and cells were sonicated. The lysate was cleared by centrifugation and loaded on a 5ml GSTrap FF column (GE Healthcare). XIAP was cleaved from the GST-tag with Prescission enzyme in 5ml of 50mM HEPES, 50mM NaCl, pH7.4 overnight at 4°C. Eluted fractions of XIAP were pooled and loaded on a 6ml Resource Q (ion exchange) column and eluted with a linear gradient of 50-500mM NaCl (50μM HEPES, pH7.4). Lastly, GE gel filtration column Superdex 200 (16/60) was used with elution buffer: 150mM NaCl, 50mM HEPES, pH 7.4. Fractions containing purified XIAP were pooled and concentrated to 1μg/µl.

### GST-Pull down assay

A method for GST-pull down assay is described elsewhere (Ikeda et al., 2011). Briefly, HEK293T were transfected with the respective constructs using GeneJuice (VWR International, #70967-3) according to the manufacturer’s recommendations. Cells were lysed in 600μl lysis buffer (50mM HEPES, 150mM NaCl, 1mM EDTA, 1mM EGTA, 25mM NaF, 10mM ZnCl2, 10% glycerol, 1% Triton-X-100, supplemented with cOmplete mini proteinase inhibitor), centrifuged and lysate was incubated with GST-proteins attached to Glutathione Sepharose 4B agarose beads overnight at 4°C. The beads were washed three times with lysis buffer and samples were separated by SDS-PAGE and analyzed by immunoblotting. Purified XIAP was diluted to a concentration of 11nM in PD buffer (150mM NaCl, 50mM Tris pH7.5, 5mM DTT, 0.1% NP-40) ^recipe provided by David Komander^. 500ng XIAP were pulled down on immobilized GST-proteins attached to Glutathione Sepharose 4B agarose beads for 4h at 4°C. The beads were washed four times with PD buffer and samples were separated by SDS-PAGE and analyzed by immunoblotting.

### Peptide competition assay

Recombinant XIAP, GST-empty and GST-GABARARAP (WT, Y49A/L50A or R67A) were used. Per reaction 60 µg of XIAP (0.2 mg/ml) were incubated with 30 µg of GST or GST-GABARAP WT or mutants (0.1 mg/ml each) in pulldown buffer (25 mM Tris, 150 mM NaCl, 10% glycerol, 0.1% Igepal, 1 mM EDTA, cOmpleteTM Protease Inhibitor Cocktail) for 2 h at 4°C. This was followed by incubation for 1h at 4°C with 20 µg of Pierce Glutathione Magnetic Beads (Thermo Scientific) per reaction. Subsequently, the beads were washed six times with pulldown buffer. Subsequently, samples in 2x Laemmli buffer containing 5 mM β-mercaptoethanol (Bio-Rad) were loaded on SDS-PAGE gels, stained by Coomassie brilliant blue and imaged on ChemiDoc MP (Bio-Rad). For the in vitro competition assay, 0.5 mM of GIM-containing peptide EDEWVNVQY or GIM mutant peptide EDEAVNAQY (each synthesized in-house) were added to the reaction mixture and proceeded as described.

### RNA sequencing

To prepare RNA-Sequencing samples, total RNA was isolated from *Xiap*^+/+^, *Xiap*^ΔRING/ΔRING^ and *Xiap*^−/−^ MEFs, grown in full medium or starved for 2h using TRIzol (Life Technologies, 15596026). DNA was digested by TURBO DNA-free Kit (Ambion, AM1907) and Bioanalyzer 2100 (Agilent Technologies) was used to determine the quality and quantity of RNA according to the manufacturer’s instructions. The library was prepared from these samples by poly(A) enrichment (New England Biolabs, Ipswich, MA). The resulting fragmented samples were sequenced on a HiSeqV4 SR50 with a read length of 50 (by VBCF-NGS). The reads were mapped to the *Mus musculus* mm10 reference genome with STAR (version 2.4.0d) (Dobin, Davis et al., 2013). Reads aligning to rRNA sequences were filtered out prior to mapping. The read counts for each gene were detected using HTSeq (version 0.5.4p3) (Anders, Pyl et al., 2015). The counts were normalized using the TMM normalization from edgeR package in R. Prior to statistical testing the data was voom transformed and then the differential expression between the sample groups was calculated with limma package in R. The functional analysis was done using the topGO and gage packages in R. For visualization, heat maps were created using R and fragment alignments were processed using the Integrative Genomics Viewer (IGV_2.3.40 software) (Robinson, Thorvaldsdottir et al., 2011, Thorvaldsdottir, Robinson et al., 2013).

### Statistical analysis

Graphs were created using GraphPad Prism 7 software (GraphPad Software, Inc). FACS data was additionally visualized as histograms using FlowJo 10.0.8r1 software. Students *t*-test was used to compare two groups. In figures, asterisks denote statistical significance as calculated by Student’s t-test (* p<0.05, ** p<0.01, *** p<0.001). RNA-Seq data were analyzed using a two-way analysis of variance (ANOVA) multiple comparison.

## Acknowledgements

We acknowledge technical supports by Franziska Christina Hofer (IMBA, Vienna, Austria) and from IMBA/IMP core facilities (Vienna, Austria) of BioOptics, bioinformatics (Maria Novatchkova and Alex Schleiffer), peptide production (Mathias Madalinski) and molecular biology service, and VBCF services (Vienna, Austria) for the next generation sequencing and electron microscopy. We also acknowledge Felix Randow, Philipp Jost, Ulrike Resch and Lynn Wong for kind gift of materials used in this study. This study was supported by the ERC consolidator grant (LUbi), the FWF standalone grant, and Austrian Academy of Sciences. We thank Life Science Editors for the support in editing this manuscript.

## Author contributions

I.P., P.E., L.D., I.B., M.S., C.D.G., Y.D. and F.I. carried out the experiments and analyzed the data. Y.D. and F.I. planned the experiments, conducted the project, and interpreted the data. F.I. designed the experiments and wrote the manuscript.

## Materials & Correspondence

Correspondence to: Fumiyo Ikeda

## Supplementary Figure legends

**Supplementary Figure 1.**
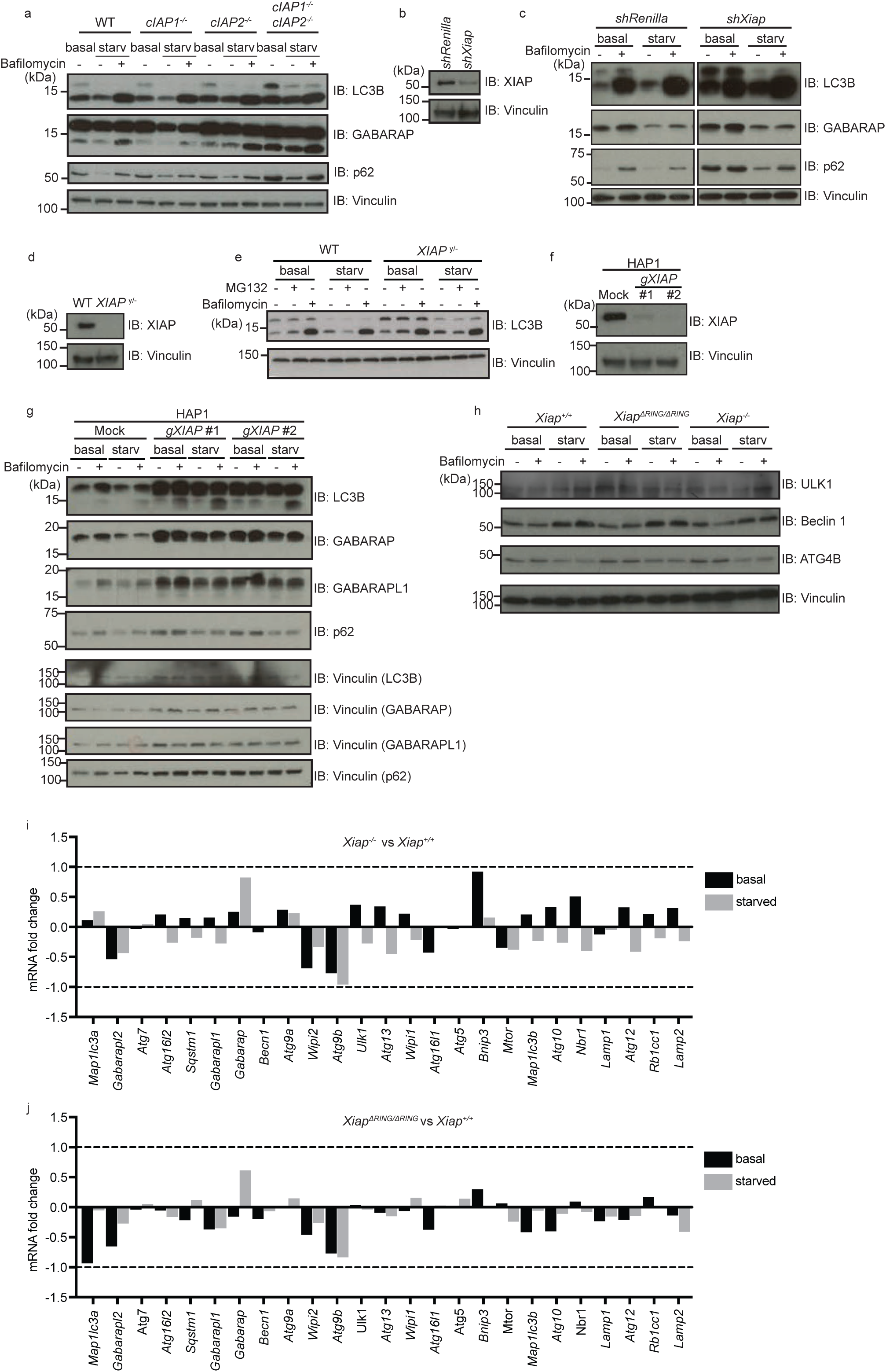
Expression levels of autophagy related proteins or genes in IAP- or XIAP-deficient cells. **(a)** Levels of autophagy-related proteins in total cell extracts of wild type (WT), cIAP1 knockout (*cIap1*^−/−^), cIAP2 knockout (*cIap2*^−/−^) and cIAP1/cIAP2 double knockout (*cIap1*^−/−^; *cIap2*^−/−^) MEFs analyzed by immunoblotting. Protein levels of LC3B, GABARAP, p62, XIAP were detected with the antibodies as indicated in MEFs in basal or starved conditions with or without Bafilomycin A1 treatment (100nM, 2 hr). Vinculin was monitored as loading control. **(b and c)** Levels of autophagy-related proteins (LC3B, GABARAP and p62) (c) and knockdown efficiency of XIAP (b) in total cell extracts of control (*shRenilla*) and XIAP-knockdown (*shXiap*) MEFs examined as (a). **(d and e)** Levels of LC3B in control (WT) and XIAP-deficient (*XIAP*^*y/-*^) HCT116 cells were examined in basal or starved conditions with or without MG132 or Bafilomycin A1. XIAP deficiency was confirmed by using anti-XIAP antibody (d) and loading was examined by anti-Vinculin antibody. **(f and g)** Levels of autophagy-related proteins (LC3B, GABARAP, GABARAPL1 and p62) were examined in CRISPR-based XIAP knockout HAP1 (human haploid) cells under different conditions as (c). Knockout of XIAP and loading were confirmed by using anti-XIAP antibody (f) and by an anti-Vinculin antibody (f and g), respectively. **(h)** Protein levels of autophagy regulators important for the initiation step (ULK1, Beclin 1 and ATG4B) were determined by immunoblot in *Xiap*^*+/+*^, *Xiap*^*∆RING/∆RING*^ and *Xiap*^−/−^ MEFs in basal or starved condition. **(i and j)** RNA-Seq data of genes involved in autophagy regulation in *Xiap*^−/−^ (i) or *Xiap*^*∆RING/∆RING*^ (j) MEFs compared to WT MEFs in basal and starved conditions. Fold change of poly (A)-enriched total RNA in *Xiap*^−/−^, *Xiap*^*∆RING/∆RING*^ and *Xiap*^*+/+*^ MEFs was compared between each other.

**Supplementary Figure 2.**
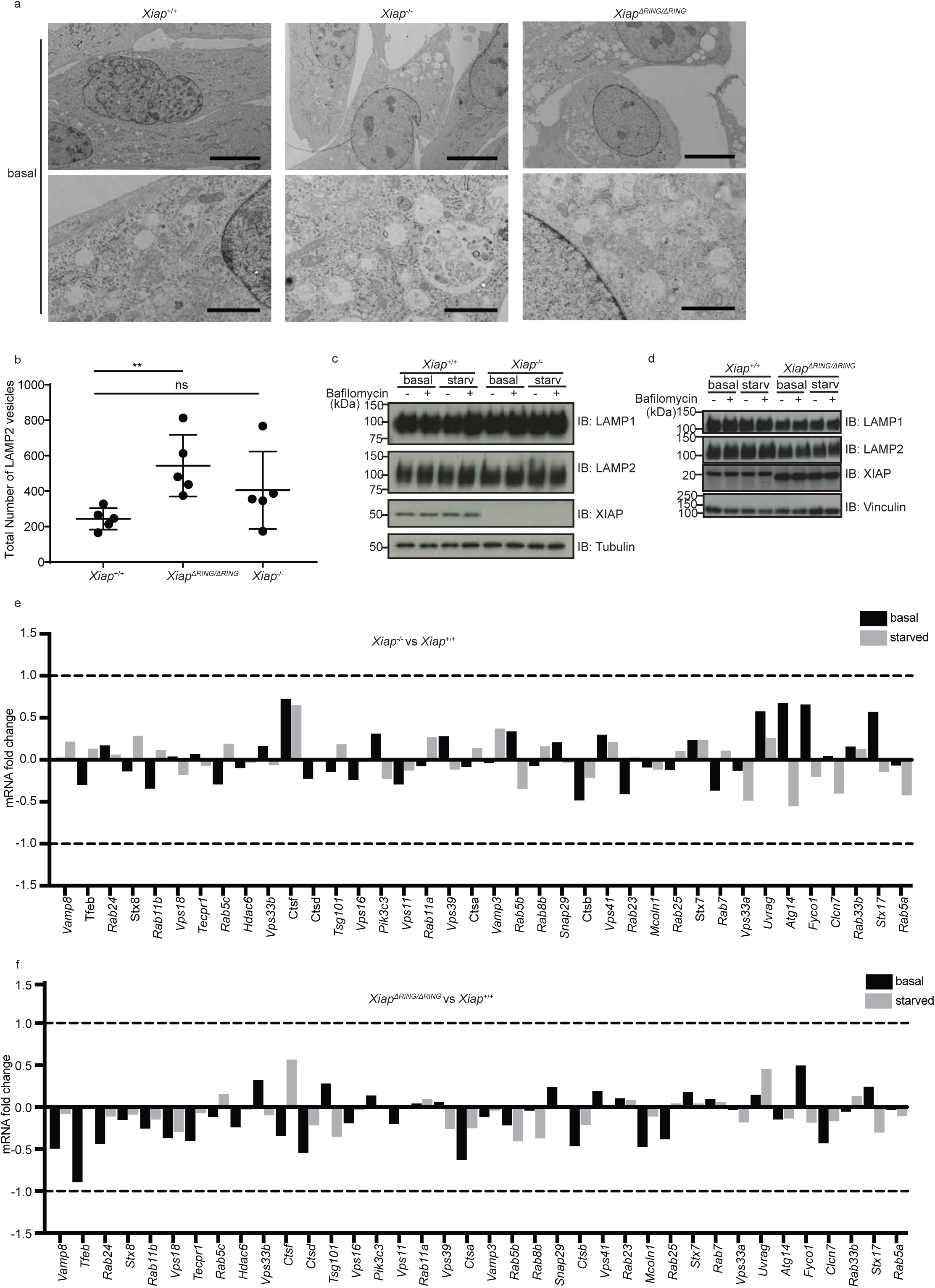
Characterization of lysosomes in *Xiap*^+/+^, *Xiap*^−/−^ and *Xiap*^*∆RING/∆RING*^ MEFs. **(a)** Electron microscopic images of *Xiap*^+/+^, *Xiap*^−/−^ and *Xiap*^*∆RING/∆RING*^ MEFs in basal condition. Scale bars, overview (top panels), 10 µm, magnified view (bottom panels), 2 µm. **(b)** Total number of LAMP2-positive structures in *Xiap*^+/+^, *Xiap*^−/−^ and *Xiap*^*∆RING/∆RING*^ MEFs under starvation treated with Bafilomycin A1 as in Figure 2d. **(c and d)** Immunoblot of *Xiap*^+/+^, *Xiap*^−/−^ (c) and *Xiap*^*∆RING/∆RING*^ (d) MEFs examining lysosome proteins LAMP1 and LAMP2. Expression of XIAP and loading was examined by using anti-XIAP, and anti-Tubulin or anti-Vinculin antibodies, respectively. **(e and f)** mRNA fold changes of lysosome-related genes in *Xiap*^−/−^ and *Xiap*^*∆RING/∆RING*^ MEFs in comparison to WT MEFs determined by RNA-sequencing.

**Supplementary Figure 3.**
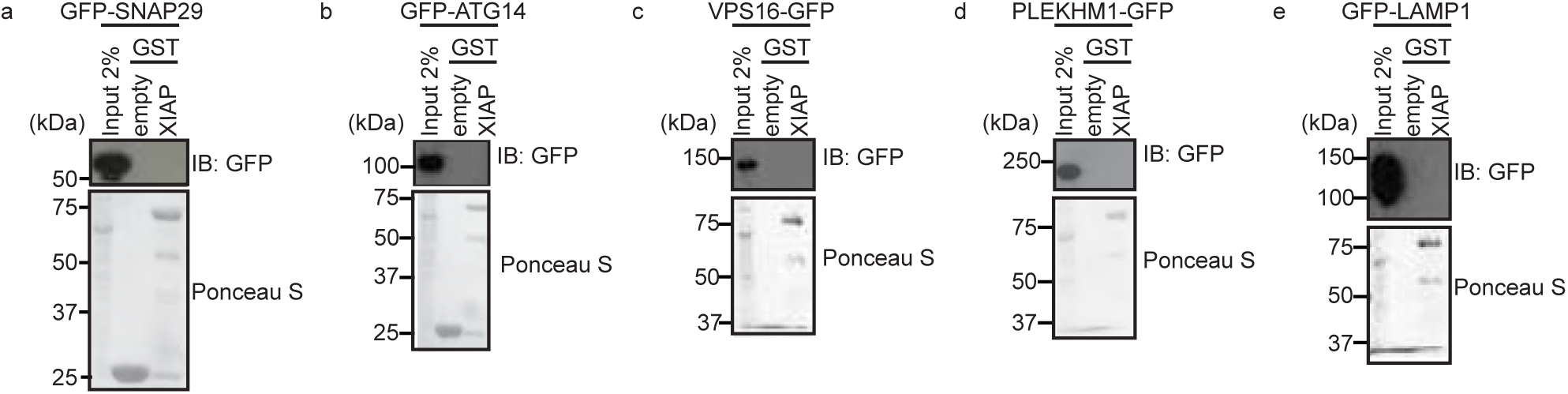
Interaction between XIAP and known autophagosome-lysosome fusion proteins. **(a-e)** Interaction between known autophagosome-lysosome fusion proteins and XIAP examined by GST-pulldown assays. Total cell extracts of HEK293T cells transiently expressing GFP-fusion proteins as indicated were incubated with Glutathione sepharose beads-immobilized GST-XIAP, pulled down, washed and examined by SDS-PAGE followed by immunoblotting using an anti-GFP antibody. GST-empty was used as control and loading of GST-fusion proteins was examined by Ponceau S staining.

**Supplementary Figure 4.**
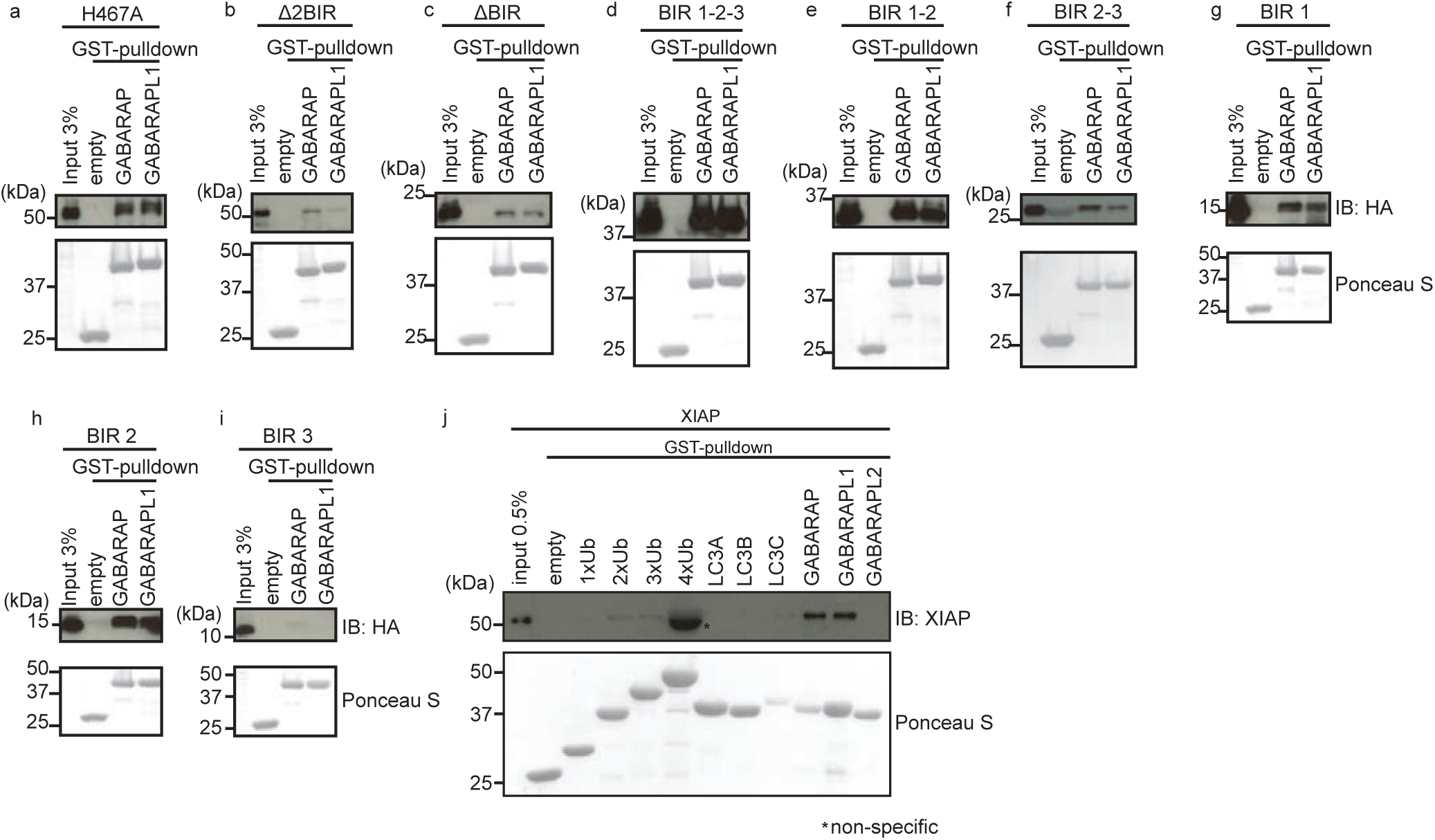
XIAP interacts with GABARAP via multiple binding sites. **(a-i)** Interaction of various XIAP mutants (as in Figure 4a) with GST-GABARAP or GST-GABARAPL1 examined by GST-pulldown assays. Total cell extracts of HEK293T cells transiently expressing HA-tagged XIAP mutants as indicated were incubated with GST-empty, GST-GABARAP, or GST-GABARAPL1 proteins immobilized with glutathione sepharose beads. Interaction was detected by immunoblotting using anti-HA antibody and loading of GST proteins were examined by Ponceau S staining. **(j)** Direct interaction between XIAP and LC3 family members or ubiquitins in different lengths examined by GST-pulldown assays similar to Figure 4b. Recombinant XIAP protein purified from *E. coli* was used instead.

